# Regional and network-level resting state functional connectivity markers of pharmacological treatment resistance in obsessive-compulsive disorder

**DOI:** 10.64898/2026.09.23.753814

**Authors:** Harleen Chhabra, Srinivas Balachander, Vani Holebasavanahalli Thimmashetty, Venkataram Shivakumar, Lekhansh Shukla, Ganesan Venkatasubramanian, YC Janardhan Reddy, Janardhanan C Narayanaswamy

## Abstract

Treatment resistance in obsessive-compulsive disorder (OCD) is common, with 40–60% of patients failing to respond to pharmacotherapy. Identifying functional differences between responders and non-responders may guide alternative treatment strategies. We investigated resting-state brain differences between selective serotonin reuptake inhibitor (SSRI) responders and non-responders and compared them with healthy controls (HC). 3T MRI scans were acquired from 80 participants with OCD and 50 HCs. Structural and 5-minute resting-state data were processed using the CONN toolbox. Regions showing group differences in amplitude of low-frequency fluctuation (ALFF) were identified for seed-based connectivity analysis. ANCOVA revealed significant ALFF differences among groups in the anterior middle temporal gyrus (R-aMTG) and temporal pole (R-TP). SSRI responders showed reductions in seed-to-voxel connectivity than non-responders between these regions and the left superior and inferior occipital cortices, left superior parietal lobule, and left temporo-occipital middle temporal gyrus. Responders also showed reductions in connectivity with the left superior parietal lobule compared with HCs. Across the OCD group, Y-BOCS scores positively correlated with ALFF in the R-aMTG (p=0.008) and R-TP (p=0.044). Altered temporal cortical activity and connectivity with posterior regions were associated with SSRI treatment response, supporting a role for temporal–posterior cortical networks in OCD treatment response.

## 1. Introduction

Obsessive compulsive disorder (OCD) is associated with significant dysfunction and disability (Fineberg et al., 2020) and high public health burden (Hollander et al., 2016). Selective serotonin reuptake inhibitors (SSRIs) and cognitive behavior therapy (CBT) are first-line treatments (Arumugham et al., 2026; Van Ameringen et al., 2026). However, a substantial proportion of patients are resistant to SSRIs (Pallanti et al., 2002). Treatment resistance is defined as non-response to one or more SSRI treatment trials and the term treatment refractory is used to indicate poor response to most of the available treatment options, including CBT and pharmacological augmentation strategies (Fineberg et al., 2015; Pallanti et al., 2004).

Despite its prevalence, the neurobiological basis of pharmacological treatment resistance in OCD remains poorly understood. MRI studies have identified structural and functional correlates of SSRI response. Pre-treatment activation of the right cerebellum and left superior temporal gyrus (STG) during symptom provocation was associated with treatment response(Sanematsu et al., 2010), while non-responders showed greater raphe nucleus–left middle temporal gyrus (MTG) connectivity than responders (Kim et al., 2019). Differences in orbito-frontal cortex (OFC) thickness (Hoexter et al., 2015) and cortical thickness and structural covariance patterns have also been associated with treatment response, with the latter predicting response with 89.0% accuracy (Yun et al., 2015). Cerebral-cerebellar connectivity was reported to normalize following 4-week SSRI treatment (Yan et al., 2024a) and resting-state connectivity of the dorsal attention and fronto-parietal networks was associated with SSRI response (Bakay et al., 2024).

Resting-state functional MRI allows examination of intrinsic brain activity without a specific task. ALFF (0.01–0.08 Hz) measures spontaneous regional brain activity and is associated with regional neuronal activity (Duff et al., 2008; Fox and Raichle, 2007). Alterations in ALFF/fALFF have been reported in OCD (Fan et al., 2017; Meng et al., 2018; Yang et al., 2019), however its relationship with SSRI treatment response remains insufficiently understood.

In the present study, therefore examined ALFF differences among SSRI responders, non-responders, and healthy controls, followed by exploratory seed-to-voxel functional connectivity analysis of regions showing significant ALFF differences. We hypothesized that responders and non-responders would differ in regional intrinsic brain activity and corresponding functional connectivity.

## 2. Methodology

### 2.1 Study participants

The sample consisted of 50 healthy volunteers (HC) (M:F= 23:27) and 80 patients (40 Non-responders, 40 Responders) (M:F= 34:46) with a diagnosis of OCD as per DSM-IV-TR criteria (Diagnostic and Statistical Manual for Mental Disorders, American Psychiatric Association). The patients were recruited from the specialty OCD Clinic of the National Institute of Mental Health & Neurosciences (NIMHANS), Bengaluru, India. A diagnosis of OCD was established after a detailed clinical interview and confirmed by using the Mini International Neuropsychiatric Interview Plus 5.0 (Sheehan et al., 1998). Handedness was assessed using the Edinburgh Handedness Inventory (Oldfield, 1971). Depressive and anxiety symptoms were assessed using the Hamilton rating scales for depression and anxiety (HAM-A and HAM-D) (Hamilton, 1959, 1960) and the obsessive-compulsive symptoms were assessed using the Yale-Brown Obsessive-Compulsive Scale (YBOCS) (Goodman et al., 1989). Participants with OCD were on a stable dose of medications for at least 3 months. SSRI responders had ≥35% reduction in YBOCS-total score and had a CGI-S score of 1 or 2 after an adequate trial with any of the SSRIs, and SRI non-responders had <25% reduction in YBOCS-total score and a CGI score > 3 after at least 2 adequate SRI trials (Shetti et al., 2005). Partial responders, that is YBOCS scores between 25 to 35, were not included in the study. The total fluoxetine equivalent dose of SSRI for non-responders and responders were 88.37±38.90 mg/day and 69.96±30.03 mg/day respectively. Other than SSRIs, twenty patients were on clonazepam (eleven non-responders and 9 responders), seven patients were on antipsychotics (six non-responders and one responder), one non-responder was on thyroxine, and one was on ondansetron. The study was carried out in accordance with the Declaration of Helsinki after obtaining institute ethical approval. All participants gave written informed consent before the assessments.

#### Resting State fMRI acquisition

MRI scans were acquired from a 3.0 Tesla MRI scanner (Magnetom Skyra). T1-weighted MRI scans were acquired for co-registration with functional images using the following parameters: TR = 8.1 msec, TE = 3.7 msec, nutation angle = 8-degree, FOV = 256 mm, slice thickness = 1 mm without inter-slice gap, NEX = 1, matrix = 256×256, yielding 165 sagittal slices. For functional scanning during rest, an Echo Planar Imaging (EPI) acquisition was obtained using a 32-channel coil for 5 minutes 14 seconds, yielding 153 dynamic scans. Following were the parameters used for acquisition: TR= 2000 msec; TE = 30 msec; flip angle = 78; Slice thickness = 3mm; Slice order: Descending; Slice number = 37; Gap = 25%; Matrix = 64 × 64 × 64 mm^3^, FOV = 192×192, voxel = 3.0 mm, isotropic.

### 2.2 fMRI data analysis

Data was acquired for a total of 50 HC and 40 each of SRI responders and non-responders. Data processing and analysis was done using Conn toolbox version 19c (Whitfield-Gabrieli and Nieto-Castanon, 2012).

First, the functional data were realigned using SPM realign & unwarp procedure (Andersson et al., 2001), where all scans were coregistered to a reference image (first scan of the first session) using a least squares approach and a 6 parameter (rigid body) transformation (Friston et al., 1995), and resampled using b-spline interpolation to correct for motion and magnetic susceptibility interactions. Potential outlier scans were identified using ART as acquisitions with framewise displacement above 0.9 mm or global BOLD signal changes above 5 standard deviations (Power et al., 2014), and a reference BOLD image was computed for each subject by averaging all scans excluding outliers. Then, functional and anatomical data were co-registered using SPM inter-modality co-registration procedure (Ashburner and Friston, 1997) with a normalized mutual information objective function. Functional data were smoothed using spatial convolution with a Gaussian kernel of 6 mm full width half maximum (FWHM). Last, anatomical data were segmented into grey matter, white matter, and CSF tissue classes using SPM unified segmentation and normalization algorithm (Ashburner, 2007; Ashburner and Friston, 2005) with the default IXI-549 tissue probability map template. In addition, functional data were denoised using a standard denoising pipeline including the regression of potential confounding effects characterized by filtered white matter timeseries, filtered CSF timeseries, motion parameters and their first order derivatives (Friston et al., 1996), followed by bandpass frequency filtering of the BOLD timeseries (Hallquist et al., 2013) between 0.01 Hz and 0.08 Hz. Following de-noising, four HC data were discarded (two with max motion > 3 SD; one with invalid scans > 30% and one with global signal change > 3 SD. Similarly, four responders and two non-responders were excluded due to max motion > 3 SD and one non-responder had invalid scans > 30%.

Amplitude of low frequency fluctuation (ALFF) and seed-based correlation analysis were performed on the remaining data (46 HCs, 36 SSRI responders and 37 SSRI non-responders). ALFF at each voxel represented the root-mean-square of BOLD signal amplitude after band-pass filtration (Yang et al., 2007). The ALFF analysis was performed without predefined anatomical regions of interest. Clusters showing significant between-group differences in ALFF were subsequently used as seeds for exploratory seed-to-voxel functional connectivity analyses.

### 2.4 Analysis Scheme

A one-way ANCOVA was conducted to compare ALFF among three groups-healthy controls (HC), SRI responders, and SRI non-responders with age and sex as covariates. The clusters showing significant between-group differences in ALFF were subsequently used as seeds for seed-to-voxel connectivity analyses. Group-wise differences in connectivity were assessed using independent-samples t-tests with age and sex as covariates, comparing HC versus SRI responders, HC versus SRI non-responders, and SRI responders versus SRI non-responders. These analyses were performed using the CONN toolbox. As an exploratory analysis, beta values from the significant ALFF clusters were extracted for all three groups and subjected to post-hoc group-wise comparisons using R software. Finally, within the OCD group, exploratory correlation analyses were conducted between ALFF and duration of illness and current Y-BOCS scores to further characterize whether the observed ALFF abnormalities were associated with illness chronicity or current symptom severity.

## 3. Results

The three groups did not differ in age (p= 0.23), sex (p= 0.96), were comparable on total duration of illness (p= 0.10), HAM-A scores (p= 0.06), and HAM-D (p= 0.55) scores. Non-responders had significantly greater current YBOCS scores (p < 0.001) and Fluoxetine equivalence dose (p= 0.027) (**Table 1**).

**Table 1:** Demographics and clinical details: OCD responders, non-responders and HC.

| Descriptive | Healthy controls<br>(n=46) | OCD non-responders (n=37) | OCD responders<br>(n=36) | Statistics | p |
| --- | --- | --- | --- | --- | --- |
|  | Mean±SD | Mean±SD | Mean±SD |  |  |
| Age (years) | 29.11±5.96 | 31.00±6.88 | 28.42±7.28 | F= 1.50 | 0.23 |
| Sex (M:F) | 21:25 | 16:21 | 15:21 | $\chi^2= 0.14$ | 0.96 |
| YoE | 13.93±4.32 | 12.81±3.83 | 12.44±3.52 | F= 1.62 | 0.20 |
| DOI (months) | - | 103.24±62.65 | 78.33±63.123 | t= 1.69 | 0.10 |
| HAM-A | - | 5.74±3.35 | 7.89±5.85 | t=-1.92 | 0.06 |
| HAM-D | - | 6.16±3.03 | 5.72±3.27 | t= 0.60 | 0.55 |
| Current Y-BOCS | - | 25.59±7.98 | 13.06±8.56 | t= 6.48 | **<0.001** |
| Fluox Equi<br>(mg/day) | - | 88.37±38.90 | 69.96±30.03 | t=2.26 | **0.027** |

SRI Responder: $\geq 35\%$ reduction in YBOCS-total score and CGI-I 1 or 2 with any SRI treatment; SRI non-responder: $< 25\%$ reduction in YBOCS-total score, CGI-I $> 3$ after at least 2 adequate SRI trials; YOE\_ Years of Educations; DOI: Duration of Illness; HAM-A: Hamilton Anxiety Rating Scale; HAM=D: Hamilton Depression Rating scale; Y-BOCS: Yale Brown Obsessive Compulsive Scale; Flox Equi: Fluoxetine SSRI equivalent dose

ANCOVA analysis for ALFF showed right anterior middle temporal gyrus (R-aMTG: 52 04 −32) and right temporal pole (R-TP: 48 22 −30) to be significantly different among the three groups. There was a significant lower ALFF at R-aMTG (Beta = −0.17) and higher ALFF at R-TP (Beta = 0.15) (**Table 2, Figure 1**). Post-hoc analysis with FDR correction and age and sex as covariates, showed that at both R-aMTG and R-TP, non-responders had significantly greater ALFF compared to both responders (R-aMTG: Mean beta = 0.31, p=0.002; R-TP: Mean beta = 0.34, p=0.002) and HC (R-aMTG: Mean beta = 0.40, p<0.001; R-TP: Mean beta = 0.41, p<0.001) (**Table 2)**.

**Table 2:**
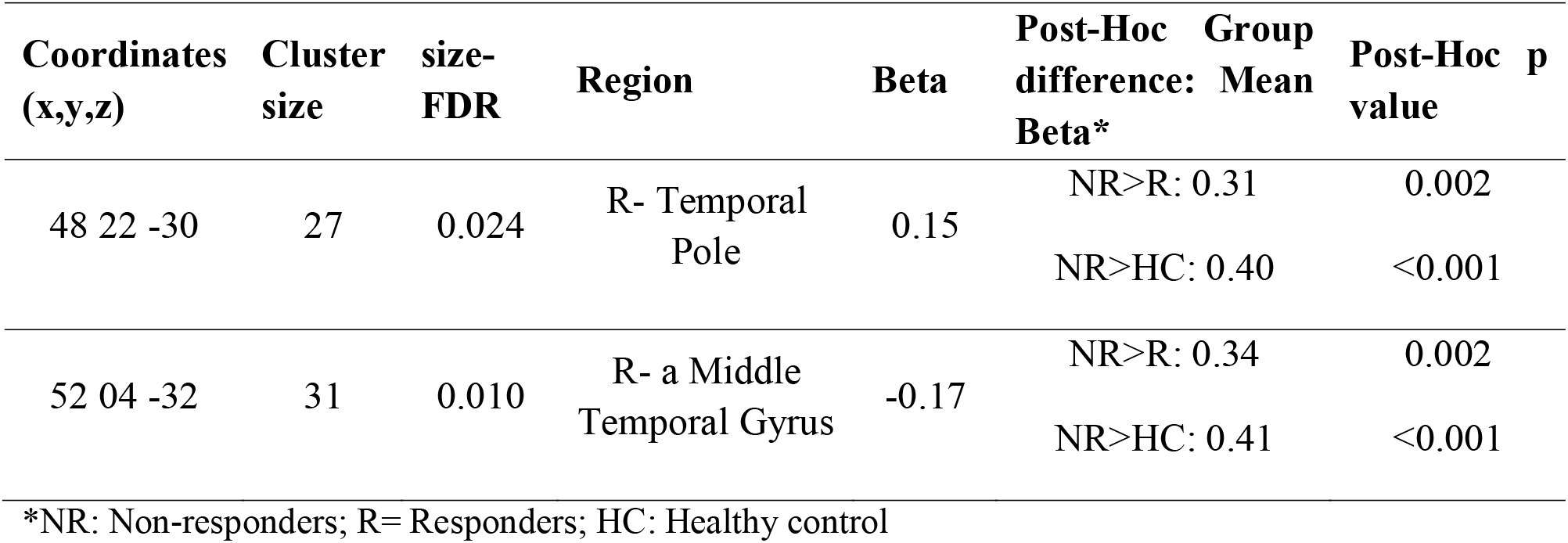
Regions with difference in ALFF between OCD responders, non-responders and HC.

**Fig 1.**
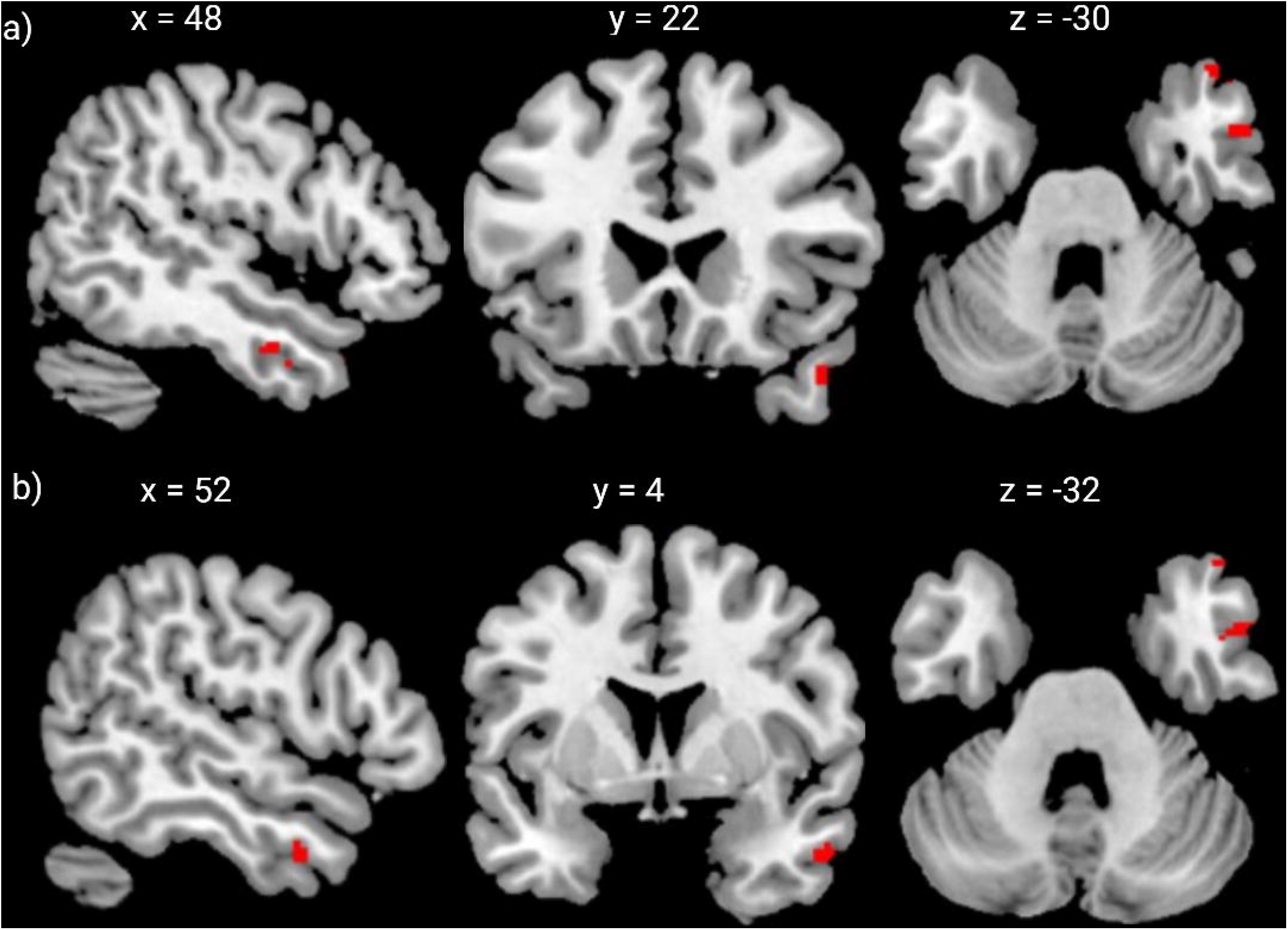
ANCOVA group differences in amplitude of low-frequency fluctuations (ALFF) across OCD non-responders, responders, and healthy controls. a) shows the cluster in the right temporal pole, b) shows the cluster in the right anterior middle temporal gyrus. Post-hoc pairwise comparisons showed significantly higher mean beta values in non-Responders compared to both responders and healthy controls for both right temporal pole and right anterior middle temporal gyrus. Statistical maps on the MNI space show thresholded results at p_FDR_ < 0.05

Given that the initial ALFF analysis did not involve predefined anatomical ROIs, the significant ALFF clusters identified in this analysis (R-aMTG and R-TP) were therefore used post hoc as seeds for an exploratory seed-to-voxel connectivity analysis.

Independent samples t-tests revealed significantly lower seed-to-voxel connectivity in SSRI responders compared with non-responders. Specifically, responders showed lower connectivity between the R-aMTG and R-TP seeds and the left superior and inferior lateral occipital cortex (cluster: −22, −60, 66; β = −0.023; cluster size = 114; size-FDR < 0.001), as well as the left inferior lateral occipital cortex (cluster: −52, −66, −02; β = −0.027; cluster size = 64; size-FDR = 0.003) (**Table 3, Figure 2**).

**Table 3:** Regions with difference in seed to voxel connectivity between OCD responders, non-responders and HC (L: left hemisphere, i: inferior, s: superior)

| **Responders > HC** |  |  |  |  |
| --- | --- | --- | --- | --- |
| **Coordinate**<br>**(x,y,z)** | **size** | **size-FDR** | **Region** | **Beta** |
| -26 -54 54 | 43 | 0.037 | L- Superior Parietal Lobule | -0.034 |
| **Responders > Non-responders** |  |  |  |  |
| -22 -60 66 | 114 | <0.001 | L- s & i. Lateral occipital cortex, L- Superior Parietal Lobule | -0.023 |
| -52 -66 -2 | 64 | 0.003 | L- i.Lateral Occipital cortex, L- temporo-occipito MTG | -0.027 |

**Fig 2.**
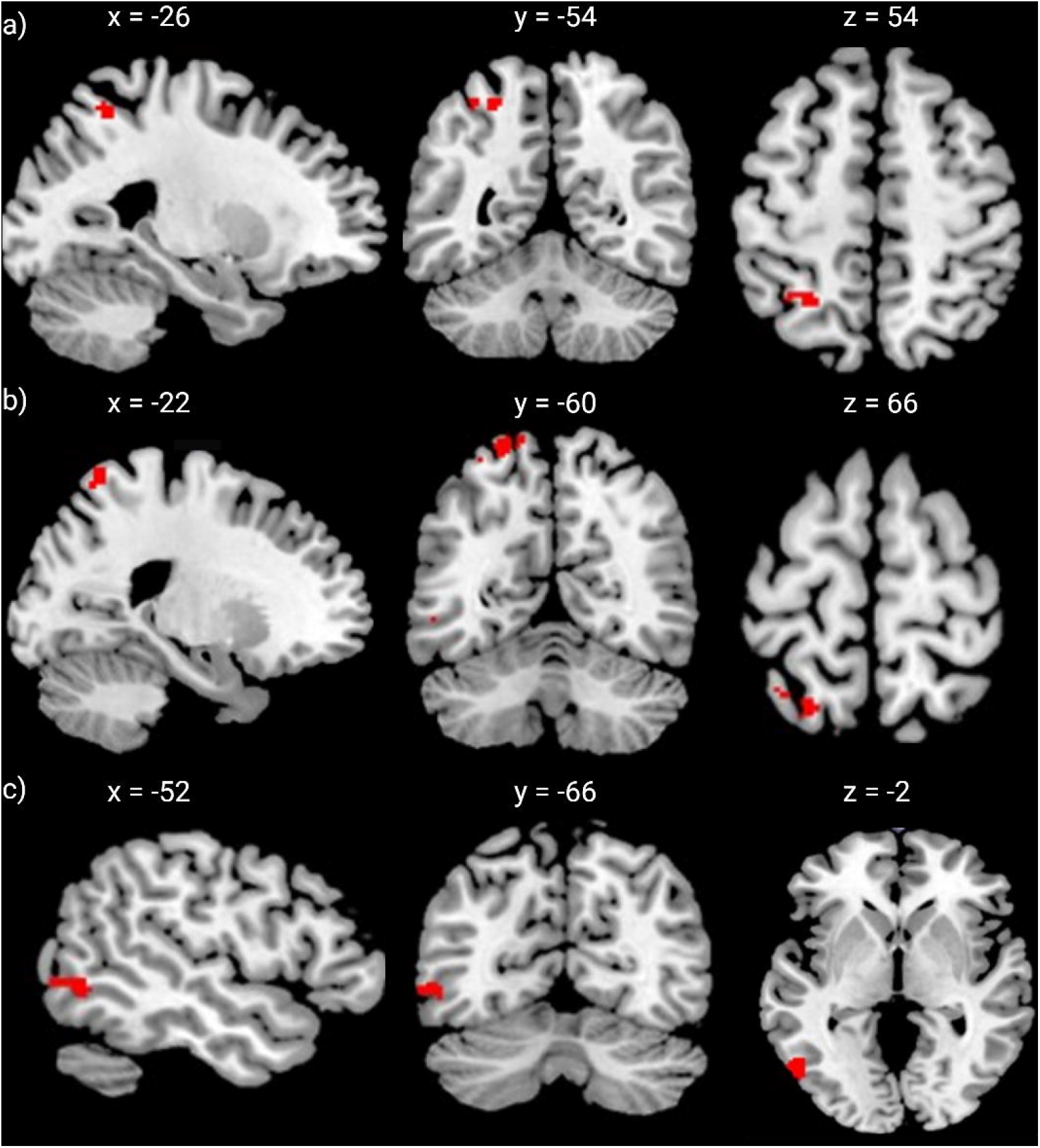
Seed-to-voxel functional connectivity differences across OCD non-responders, responders, and healthy controls. a) shows increased functional connectivity responders in the left superior parietal lobule compared to controls. b) shows increased functional connectivity in responders in the left superior/inferior lateral occipital cortex and left superior parietal lobule compared to non-responders. c) increased functional connectivity in responders in the left inferior lateral occipital cortex and left temporo-occipital middle temporal gyrus compared to non-responders. Statistical maps on the MNI space show thresholded results at p_FDR_ < 0.05

Compared with healthy controls, responders also showed significantly lower seed-to-voxel connectivity between the R-aMTG and R-TP seeds and the left superior parietal lobule (cluster: −26, −54, 54; β = ™0.034; cluster size = 43; size-FDR = 0.037). No significant connectivity differences were observed between non-responders and healthy controls (**Table 3, Figure 2**).

Exploratory correlation analyses conducted separately within the responder and non-responder groups did not reveal any significant associations between ALFF and current Y-BOCS scores. When the OCD patient group was considered as a whole, however, current Y-BOCS scores showed significant positive correlations with ALFF in both the R-aMTG (r = 0.31, p = 0.008) and R-TP (r = 0.24, p = 0.044). No significant correlations were observed between ALFF in either cluster or duration of illness.

## 4. Discussion

In the current study, using a hypothesis-independent approach, we showed differences in intrinsic brain activity and functional connectivity among SSRI responders, non-responders, and healthy controls. SSRI non-responders showed greater ALFF in the R-aMTG and R-TP than both responders and controls. Seed-to-voxel analyses based on these regions revealed lower connectivity in responders than non-responders between both temporal seeds and several posterior cortical regions. Responders also showed lower connectivity with the left superior parietal lobule than healthy controls. These findings implicate temporal-lobe activity and its connectivity with posterior cortical regions in differentiating SSRI treatment response in OCD. Together, these findings implicate temporal-lobe intrinsic activity and its connectivity with posterior cortical regions in the differentiation of SSRI treatment response in OCD.

Although OCD has traditionally been associated with abnormalities in cortico-striato-thalamo-cortical (CSTC) circuits (Jalal et al., 2023), increasing evidence indicates involvement of other cortical and subcortical regions (Jijimon et al., 2026; Milad and Rauch, 2012). Consistent with previous reports of altered ALFF/fALFF in OCD (Fan et al., 2017; Meng et al., 2018; Yang et al., 2019), our findings showed the temporal cortex, particularly the MTG and temporal pole, as regions potentially relevant to OCD and SSRI treatment response. Yang et al. reported that fALFF in the right MTG contributed to differentiating patients with OCD from healthy controls, with an overall classification accuracy of 72% (Yang et al., 2019). Other studies have also reported altered ALFF/fALFF in the MTG in OCD (Fan et al., 2017; Meng et al., 2018). Our findings extend these observations by showing that ALFF in the R-aMTG and R-TP differed according to SSRI treatment response, with higher ALFF in non-responders. Previous studies have also reported associations between temporal-lobe abnormalities and treatment response in OCD, including lower global functional connectivity density in the left MTG in non-responders compared with remitted patients (Shan et al., 2019) and involvement of temporal gyri in response to exposure and response prevention (Shi et al., 2021).

The seed-to-voxel findings are also consistent with previous evidence implicating posterior cortical regions in OCD. Altered occipital cortical function and metabolism, including changes following treatment, have been reported in previous studies (de Joode et al., 2024; Ljungberg et al., 2017; Zhao et al., 2017). The parietal cortex has also been implicated in OCD, with studies reporting altered connectivity and abnormalities related to cognitive control (Boedhoe et al., 2018; Liu et al., 2023; Liu et al., 2021; Nakao et al., 2014; van den Heuvel et al., 2016; Yan et al., 2024b). In addition, decreased functional connectivity involving occipital and temporal regions has been reported in patients with OCD compared with healthy controls and unaffected relatives (Hou et al., 2014; Tian et al., 2016), and reduced connectivity involving the middle occipital gyrus has also been reported (Jia et al., 2020). In this context, our finding of lower connectivity between the temporal seeds and lateral occipital, temporo-occipital, and superior parietal regions in responders compared with non-responders suggests that the functional relationship between temporal and posterior cortical regions may differ according to SSRI treatment response.

The observed connectivity differences may reflect differences in the interaction between temporal regions involved in emotional and belief-related processing and posterior regions involved in visual and cognitive processing. However, given the cross-sectional design of the study and the absence of pretreatment neuroimaging, it is not possible to determine whether this connectivity differences are associated with the effects of SSRI treatment or represent pre-existing differences between responders and non-responders. Rather, they suggest that altered connectivity between temporal and posterior cortical regions may be associated with SSRI treatment response in OCD. Overall, the findings extend previous evidence implicating the MTG and posterior cortical regions in OCD and suggest that their functional relationship may be relevant to treatment response.

### Strengths and limitations

This study has several strengths. Non-response to SSRI was defined stringently, including patients who had failed at least two adequate trials, and patients were on a stable medication dose for at least 3 months. Using ALFF findings to identify regions with significant group differences, followed by seed-to-voxel functional connectivity analysis, allowed examination of connectivity patterns without restricting the analysis to predefined anatomical regions, potentially identifying differences beyond the conventional CSTC. Finally, based on the observed group means and standard deviations, the estimated omnibus effect size (Cohen’s *f*) was 0.42. With a total sample of 119 participants across three groups and an alpha level of 0.05, the corresponding post-hoc power was approximately 98%.

This study has limitations. First, responder and non-responder groups differed in illness severity and mean SSRI dose (fluoxetine equivalents), which may have influenced group differences. Second, the cross-sectional design and absence of pretreatment neuroimaging preclude determining whether ALFF and connectivity differences were pre-existing markers of SSRI response or developed during treatment. Longitudinal studies with pretreatment and post-treatment imaging are needed. Third, differences in treatment response across OCD symptom dimensions were not examined, despite the clinical heterogeneity of OCD.

Our findings contribute to growing evidence of broader, diffuse neural involvement in OCD, implicating temporal lobe structures in SSRI treatment response. Differences in functional connectivity between temporal and parieto-occipital regions provide preliminary evidence of the neurobiology of medication response in OCD. Temporal lobe dysfunction and impaired communication with posterior cortical regions may represent markers of SSRI treatment resistance. However, the cross-sectional design precludes determining whether these connectivity differences preceded treatment and contribute to response or emerge during treatment. Future studies should replicate these findings in larger, longitudinal cohorts to establish the temporal stability and predictive value of temporal-parietal and temporal-occipital alterations. Multimodal imaging integrating structural, functional, and metabolic measures may further elucidate the neurobiological basis of treatment response variability. Characterizing these circuits may refine models of OCD and support more personalized treatment strategies.

## CrediT

**Harleen Chhabra:** Writing – review & editing, Writing – original draft, Visualization, Formal analysis, Data curation**. Srinivas Balachander and Lekhansh Shukla:** Writing – review & editing, Investigation, Data curation. **Vani HT:** Writing – review & editing, Investigation**. Venkataram Shivakumar:** Writing – review & editing, Visualization, Formal analysis**. Janardhanan C. Narayanaswamy:** Writing – review & editing, Supervision, Clinical Investigation, Funding acquisition, Project administration, Resources, Conceptualization, Validation. **Ganesan Venkatasubramanian and YC Janardhan Reddy:** Writing – review & editing, Validation, Supervision, Resources, Methodology, Conceptualization.

## Data availability

The data that support the findings of this study are not openly available due to reasons of sensitivity, and anonymized data will be made available from the corresponding author upon reasonable request.

